# On the relationship between serotype, chemotype and genotype for Cryptococcus spp. including a method for including polysaccharide structure in strain characterization

**DOI:** 10.1101/2024.08.22.609096

**Authors:** Maggie P. Wear, Seth D. Greengo, Scott A. McConnell, Alan Xu, Livia Liporagi Lopes, Ananya Majumdar, Arturo Casadevall

## Abstract

Over the past eight decades the classification of cryptococcal strains has relied on the technologies available to discriminate among isolates, such as serology, exopolysaccharide (EPS) NMR, and most recently multi-locus sequencing, which yielded serotypes, chemotypes, and genotypes, respectively. However, as one method superseded the other, the relationship between classification schemes became uncertain, resulting in assumptions that have not been rigorously validated. Here we compared the serotype, chemotype, and genotype as defined by multi-locus sequence typing (MLST) of 63 strains for which both serotype and chemotype characterization was available. None of the three strain typing methods were correlative with each other, although for about 50% of the strains there was correlation between chemotype and genotype. To address this, we updated the methodology of GXM motif categorization for cryptococcal strains through analyzing filter-isolated EPS by 1D [^1^H] NMR and assigning GXM motifs along with the O-acetylation level. The result is a facile method using minimally processed EPS material coupled with simple cryptococcal strain classification by GXM motif expression. While this method increases the correlation between MLST genotype and GXM motif expression, further studies to establish the polysaccharide regulatory genes in cryptococcus will be necessary to understand the polysaccharide differences. In summary, none of the current classification methods correlated with each other, indicating a dissociation between genotype and phenotype, which poses a challenge to the cryptococcal field for explaining how phenotypic characteristics arise and are maintained.

**Author Summary:** This work examines the relationship between methods of cryptococcal strain classification and the primary structure of the predominant polysaccharide glucuronoxylomannan (GXM). We find no strong correlation between the expression of trimannose repeat motifs of GXM by a strain and serotype, MLST genotype, or the previous polysaccharide classification system, chemotyping. With an understanding that none of these classification systems accurately represent the GXM sequences in cryptococcal polysaccharides, we describe a new, facile, rapid method of polysaccharide isolation and GXM motif expression characterization. Application of this method reveals a strong association between high levels of GXM O-acetylation, necessary for antibody binding, and the expression of a single motif of GXM. Additionally, the association between MLST genotype and GXM motif expression is higher when utilizing this new method. While much remains to be understood about this complex system, this work provides a new method to include GXM motif expression in cryptococcal strain expression allowing for more accurate strain descriptions in the future.

## Introduction

*Cryptococcus neoformans* and *Cryptococcus gattii* are species complexes (1) unique among fungal pathogens in that the fungal cell is encased in a thick polysaccharide (PS) capsule that is antiphagocytic. The capsule is considered to be the most important contributor to the virulence composite of *Cryptococcus* spp (2). In addition, *Cryptococcus* spp. release exopolysaccharide (EPS) into infected tissues that interfere with the host immune response (3, 4, 5, 6, 7, 8). Over the past century cryptococcal strains have been categorized using the methods available at the time, which at first consisted of serological adsorption analysis and more recently includes molecular typing of genomic sequences. However, classification based on PS has been largely ignored due to the variability of structure and the need for specialized NMR methods. As newer technologies were introduced, the various classification schemes have not been reconciled or assembled into a coherent methodology.

The first classification of cryptococcal strains was done by Benham in the 1930s and was based on antigenic differences detected by antibody agglutination techniques (9). She defined four groups wherein no cross agglutination with other members occurred. However, for at least one of these groups production of the sera required treatment of the yeast cells with hydrochloric acid to partially remove the capsule prior to rabbit inoculation (9, 10). Evans and colleagues followed up Benham’s work, immunizing rabbits with formalin killed encapsulated yeast cells for their antigenic characterization (10, 11). Using this approach three antigenic groups were identified, known as serotypes A, B, and C (11, 10). However, serology was not fully discriminatory as cross reactions were observed in both tube agglutination and serum capsular reactions. While much of the cross-agglutination effects could be removed by adsorption with the same strain, some remained. This cross-agglutination was noted by the authors in both publications (11, 10). Description of the fourth serotype in cryptococcal species did not occur until almost twenty years later when Wilson & colleagues serotyped 106 cryptococcal strains (12). With this larger collection, the authors delineated four groups including the final serotype – serotype D – although they continued to observe the same cross-agglutination Evans and colleagues did in the 1950s. In addition to cross-agglutination, some strains typed both A and D resulting in a fifth serotype delineation – serotype AD (12, 13, 14).

The next great leap in our understanding of strain differences came from NMR structure studies of cryptococcal exopolysaccharides (EPS), which began in the 1970s and continued through the 1990s. Work carried out primarily by Cherniak and Battacharjee, and their collaborators, resulted in the identification of the major EPS polymers – glucuronoxylomannan (GXM) and glucuronoxylomannogalactan (GXMGal), previously thought to be galactoxylomannan (GalXM) and the description of their primary structure (15). Even the earliest of these publications (16) asserted that antigenic differences among the polysaccharides defined the cryptococcal serotype. NMR analysis of capsular material revealed that GXM was polymer of repeating units, known as triad motifs, composed of three mannoses, a glucuronic acid and a variable number of xylose residues. By the time Cherniak and collaborators published work that utilized an artificial neural network to classify cryptococcal strains by their constituent triad motifs, most publications stated that the M1 motif was expressed by serotype D, M2 by serotype A, M3 by serotype B, and M4 by serotype C cryptococcal strains (17). Though a link between serotype classification and PS structure was often assumed, that link was never established. In that same work two further GXM motifs (M5 & M6) are reported but also not correlated with a serotype (17). The method of categorizing cryptococcal strains by GXM motif expression proposed by Cherniak, chemotyping, was never adopted by the wider cryptococcal field. This is likely due to the difficulty of the method – it is time consuming, requires specialized equipment and analysis, and assignment of the triad residues in many cases depends upon accurate disambiguation of overlapping or degenerative chemical shift assignments, resulting in uncertainty.

Concomitant with the development of chemotyping as a classification strategy, genetic characterization using specific gene sets was developed (18, 19). Several groups put forth different gene sets for strain discrimination, but eventually a consensus emerged around set of genes referred to as Multi-Locus Sequence Typing (MLST) (20). MLST is based on amplicon sequencing of genes encoding capsule polysaccharide (*CAP59*), glycerol 3-phosphate dehydrogenase, (*GPD1*), laccase (*LAC1*), phospholipase B1 (*PLB1*), superoxide dismutase (*SOD1*), orotidine monophosphate pyrophosphorylase (*URA5*) genes and the intergenic spacer (IGS1) region (20). Together the specific sequences of the listed seven genes make up the molecular type of a cryptococcal strain. Although MLST classification is based on genetic markers that are not exclusively related to polysaccharide synthesis, they are correlated with strain serotype for most but not all strains (Table 1). Further, MLST is database-supported such that a researcher can utilize PCR reference sequences to determine the molecular type of a new strain (https://mlst.mycologylab.org).

**Table 1:**
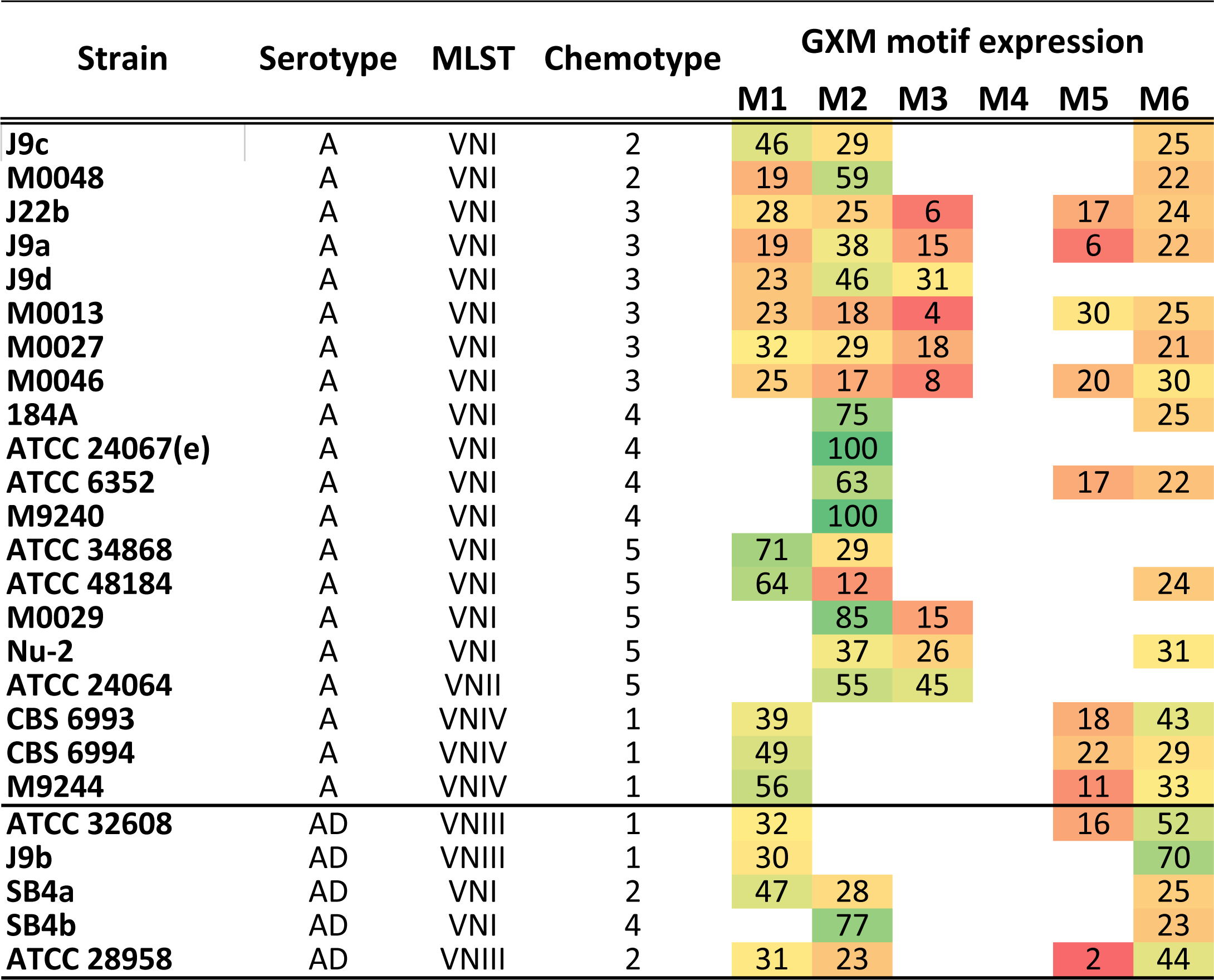
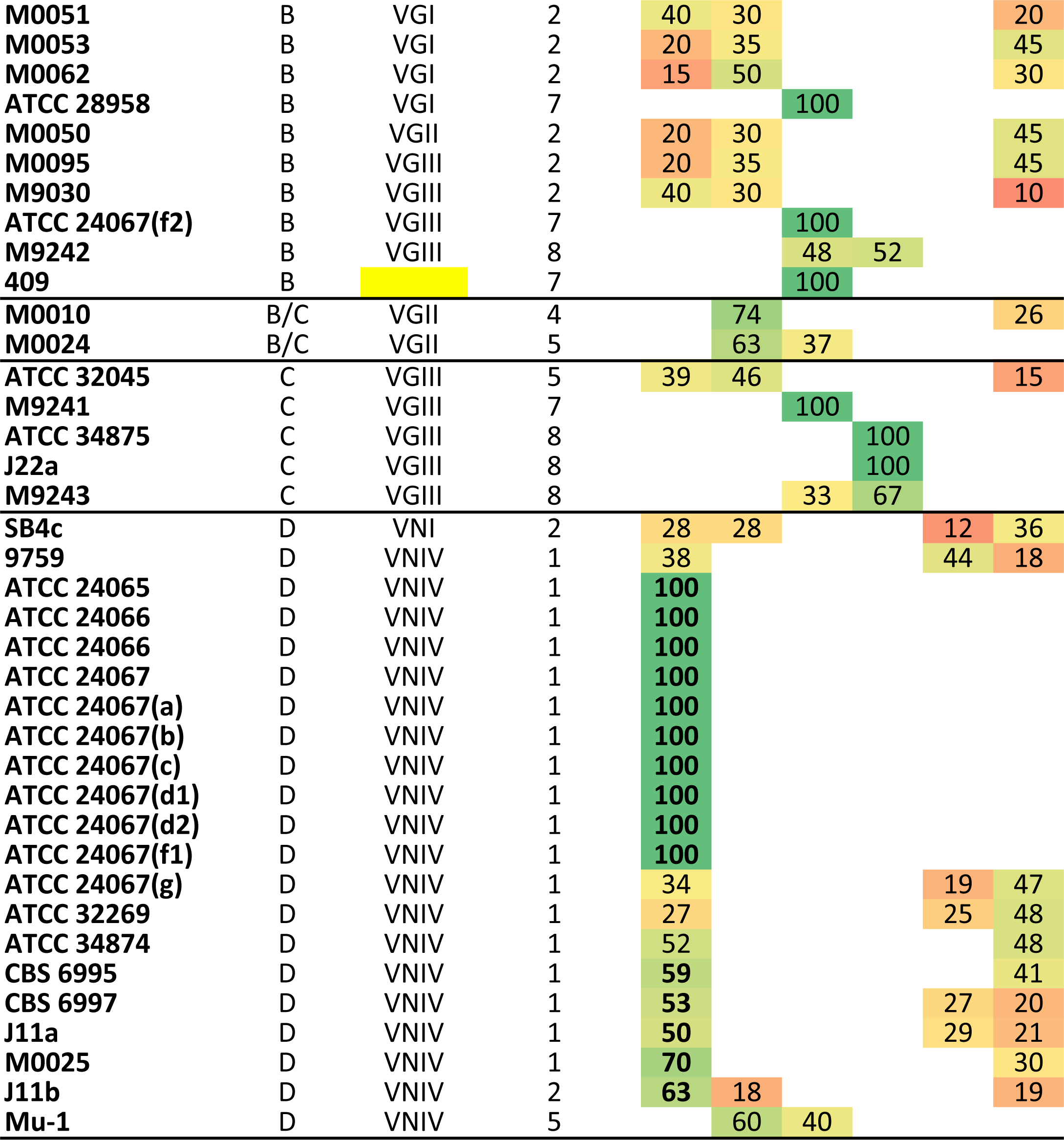
Comparison of Cryptococcal categorization technique to GXM motif expression.

In this study we examined the concordance between the three different methods of cryptococcal strain classification: serotype, chemotype, and molecular type. To do this, we collated data on as many strains as possible with respect to serotype, chemotype, and molecular type as well as data on GXM motif expression and strain source (environmental or clinical). A total of 63 cryptococcal strains with complete information for these five categories were included in this analysis.

## Methods

### Classification analysis

Surveying the literature, we amassed a collection of cryptococcal strains for which full classification information was available in four categories – serotype, source (environmental or clinical), molecular type/MLST, and GXM motif expression. We then attempted to align these four categories for each strain using the most general and oldest category, serotype. In certain cases, we conducted MLST-based molecular typing or NMR-based GXM chemotyping to generate full categorical information for important strains. This analysis is summarized in Table 1 and the full analysis can be found in Supp. Table 1.

### Methods of EPS isolation

Two methods of EPS isolation are examined in this work, CTAB and sterile filtration. The CTAB method of cryptococcal EPS isolation is well documented and results in the preferential recovery of GXM. For this study we followed the process described in (17). The filtration method was described previously (21) (22) (23). In this work we updated this method. *C. neoformans* or *gattii* strains were inoculated from frozen stock directly into YPD media and grown for 2 d at 30°C with rotation. Cells were washed with PBS and inoculated at 1 x 10^5^ cells/mL into minimal media (15 mM dextrose, 10 mM MgSO_4_, 29.3 mM KH_2_PO_4_, 13 mM glycine, 3 µM thiamine-HCl; adjusted to pH 5.5 and sterile filtered) grown for 3 days at 30°C with shaking. The culture was centrifuged (4,000 x g for 15 min.) to pellet cells, and the supernatant was sterilized by passage through a 0.22 µm filter to yield sterile whole EPS (wEPS). To remove media components and concentrate the samples, wEPS was filtered through 3 kDa Molecular Weight Cut Off (MWCO) Amicon Ultra Centrifugal Filters (Millipore #UFC9003) to yield the filtered EPS sample. Graphical representation of this isolation technique can be found in Supp. Figure 1.

### Solution NMR

Solution NMR data were collected on a Bruker Avance II (600 MHz) NMR equipped with a triple resonance TCI cryogenic probe and Z-axis pulsed field gradients. 1D [^1^H] spectra were collected at a range of temperatures from 30 - 80°C, with most samples collected at 60°C, 128 scans, and a free induction decay size of 65536 points. Standard Bruker pulse sequences were used to collect the 1D data (p3919gp and zggpw5). Data were processed in Topspin (Bruker version 3.5) by truncating the FID to 8192 points using a squared cosine bell window function and zero filling to 65536 points. Spectra were analyzed using Topspin with D_6_DSS set to [^1^H] 0.00 ppm for the identification of peak chemical shifts and an integration of 1.00 for comparison to other peak integrals. To translate the original assigned SRG peaks from the CTAB EPS spectra at 80°C to filtered EPS spectra at 60°C, both CTAB and filtered EPS sample spectra were taken at 60°C and 80°C, the peaks were matched at both temperatures and are reported here at 60°C.

Two-dimensional NMR spectra were collected at 60°C with a ^1^H spectral window of 7142.857 Hz (11 ppm) and a ^13^C window of 13893.636 Hz (92 ppm) with carrier frequencies of 4.72 ppm and 61.00 ppm in ^1^H and ^13^C, respectively, 4096 points were taken in ^1^H and 128 points in ^13^C for 1H-13C HSQC with 256 scans (Bruker pulse sequence c13_ghsqc_se_basic).

### CASPER simulation NMR

We utilized the Computer Assisted Spectrum Evaluation of Regular Polysaccharides (CASPER) online simulation software to predict the NMR chemical shifts of each of the GXM motifs M1-M6 (24). The glycan structures used for prediction of the NMR chemical shifts are as follows: M1 - β-d-GlcA^v^ (1→2) [β- d-Xyl^iv^ (1→2) α-d-Man^iii^ (1→3) α-d-Man^ii^ (1→3)] α-d-Man^i^, M2 - β-d-Xyl^vi^ (1→2) [β-d-Xyl^v^ (1→2) [β-d-GlcA^iv^ (1→2) α-d-Man^iii^ (1→3)] α-d-Man^ii^ (1→3)] α-d-Man^i^, M3 - β-d-GlcA^vii^ (1→2) [β-d-Xy^lvi^ (1→2) [β-d-Xyl^v^ (1→2) α-d-Man^iv^ (1→3)] α-d-Man^iii^ (1→3)] [β-d-Xyl^ii^ (1→4)] α-d-Man^i^, M4 - β-d-GlcAv^iii^ (1→2) [β-d-Xylv^ii^ (1→2) [β-d-Xyl^vi^ (1→2) [β-d-Xyl^v^ (1→4)] α-d-Man^iv^ (1→3)] α-d-Man^iii^ (1→3)] [β-d-Xyl^ii^ (1→4)] α-d-Man^i^, M5 - β-d-GlcAv^ii^ (1→2) [β-d-Xyl^vi^ (1→2) [β-d-Xyl^v^ (1→4)] α-d-Man^iv^ (1→3) α-d-Man^iii^ (1→3)] [β-d-Xyl^ii^ (1→4)] α-d-Man^i^, and M6 - β-d-GlcA^iv^ (1→2) [α-d-Man^iii^ (1→3) α-d-Man^ii^ (1→3)] α-d-Man^i^. We noted that predicted chemical shifts changed depending upon the direction that the polymer was built and therefore maintained the original motif structure with Man^i^ being equivalent to M_a_, with a GlcA branch. We also utilized the substituent advanced option to include O-acetylation on the 6 position of different mannose residues of the GXM motifs. The “Predict NMR chemical shifts” module of CASPER yields a table of chemical shifts for each residue and position as well as a [^1^H] [^13^C] HSQC with predicted peak locations (24).

### New EPS isolation workflow and GXM motif classification

To update the GXM motif NMR classification we first compared the chemical shifts of the 1D [^1^H] NMR spectra from the CTAB, CTAB-OAc and filtered EPS sample spectra at different temperatures ranging from 30-80°C. While aligning these 1D [^1^H] spectra and examining the peaks in the reported SRG region (5.4-5.0 ppm) we observed different numbers of signals depending on the method of preparation with CTAB having the least and filtered EPS isolation having the most. Importantly, we were able to overlay these spectra at different temperatures to translate the previous CTAB EPS GXM motif mannose anomeric chemical shifts in the ascribed SRG region at 80°C to the filtered EPS samples at 60°C. Along with the CASPER chemical shift prediction and other published cryptococcal strain CTAB EPS NMR chemical shifts, these translated chemical shifts allowed us to put forth an updated consensus chemical shift table for the anomeric residues of the GXM motifs (Figure 3).

## Results

### Correlation between methods of Cryptococcal strain classification

We were interested in understanding how each classification methodology for cryptococcal strains related to one another and which method best represent the polysaccharide features of different strains. Consequently, we collected 63 strains for which there existed information on five categorical datasets, serotype, chemotype, molecular type, GXM motif expression and source – clinical or environmental (Supp. Table 1). We present a simplified analysis table with a heatmap to illustrate the GXM motif composition for each strain (Table 1). As discussed above, Cryptococcal strains are divided into four serotypes, A, B, C, and D, although there are reports of serotype AD and B/C strains (12)(13)(14), so these four are not exclusive categorizations. MLST yields molecular types VNI-IV for *C. neoformans* strains and VGI-IV for *C. gattii* strains (20). More specifically, serotype A strains are VNI and VNII (with a few VNB which is evolutionarily very similar to VNI, for more see (25)), serotype AD strains are VNIII, and serotype D strains are VNIV while the specific delineation of molecular types for *C. gattii* strains are not clearly delineated (20). Updated classification of *C. gattii* molecular types did not utilize serotype, but rather further strain specification wherein VGI is *C. gattii*, VGII is *C. deuterogattii*, VGIII is *C. bacillisporus*, VGIV is *C. tetragattii*, and VGIIIc/VGIV is *C. decagattii* (26). Examining our dataset for a correlation between serotype and molecular type is most clear for *C. neoformans* strains. Of the twenty serotype A strains, 85% are VNI or VNII while of the nineteen serotype D strains 95% are VNIV and of the six serotype AD strains 50% are VNIII. Strain serotype and MLST classification were somewhat correlative, such that at least 50% of *C. neoformans* strain molecular type and serotype aligned.

However, this correlative trend does not hold once chemotype is introduced. Chemotypes are derived from the structural reporter group (SRG) peak chemical shifts representing the anomeric carbons of the three mannose residues in the GXM motif and delineated by the predominance of the motif expressed in majority rank. However, chemotype categories are not assigned by the most commonly expressed motif, but rather in order of preference and appearance beginning with the GXM M1 motif and do not include the contributions of GXM motifs M5 and M6. Chemotype 1 strains have the M1 motif as their primary component while Chemotype 2 expressed M1 and M2, with M1 being preferred. Chemotype 3 expresses M1, M2, and M3, but again with M1 being preferred. Chemotype 4 varied from this M1 preference as this set is defined by only M2 being expressed. This is followed by chemotype 5 which expresses both M2 and M3 with M2 being preferred and chemotype 6 which expresses M2, M3, and M4 but again with M2 being preferred. Chemotype 7 is based upon the expression of the M3 motif of GXM while the final chemotype 8 includes both M3 and M4, but with M3 being preferred (17). Although these chemotypes are derived from the chemical shifts of the anomeric carbons of the three mannose residues in the GXM motif, they did not correlate with either serotype or molecular type (Table 1, Supp Table 1). Within the twenty serotype A strains are chemotypes 1, 2, 3, 4, and 5, which correlate with the dominant GXM motif of M2 for chemotypes 4 and 5 but M1 for chemotypes 1, 2, and 3 (Supp Table 1) (17). The molecular type VNI strains span chemotypes 2, 3, 4, and 5, again showing no conservation, but interestingly the VNIV strains that are serotype A are all chemotype 1. When expanded to all VNIV strains there are just two strains that are not chemotype 1 (one is chemotype 2, the other 5). Examination of three VNIII strains shows they are chemotype 1 and 2, but the sample size is too small to conclude a trend for this molecular type. With regards to the VG molecular types and chemotype, the four VGI strains are chemotypes 2 and 7 while the three VGII strains are chemotypes 2, 4, and 5, and the nine VGIII strains are chemotypes 2, 5, 7 and 8. Unfortunately our collection does not contain any VGIV strains (Table 1). When attempting to correlate chemotype with molecular type and serotype we find that chemotype correlated with molecular type 58.2% of the time and serotype 52.4% of the time. This breaks down to 52.5% of chemotype 1, 2, and 3 strains (n=40) are VNIV and 47.5% serotype D, 77% of chemotype 4, 5, and 6 strains (n=13) are VNI or VNII and 69% serotype A, and 50% of chemotype 7 and 8 strains (n=8) are serotype B (Table 1).

In addition, we observed that within this collection there were three sets of strains that were named differently but were, in fact, the same strain (Supp. Table 2). Two of the three were conserved for serotype and molecular type, and one for chemotype, but all showed variation in the GXM motifs expressed (Supp. Table 2). This is consistent with PS structural microheterogeneity over time in culture as previously reported in one of the sets of strains, 24067 (27).

### Comparison of filtration and CTAB EPS isolation by NMR

With the observation that cryptococcal strain categorization by chemotype did not correlate with serotype or molecular type, we were interested to further examine the method of polysaccharide isolation and analysis used to determine chemotype. Serotype was determined based on the antigenicity of the surface of the cryptococcal capsule while chemotype utilized secreted EPS for characterization. Capsular polysaccharide (CPS) and EPS exhibit important biophysical differences (22), although compositional differences at the molecular level have yet to be determined. Additionally, chemotyping analysis relied upon CTAB precipitation and lyophilization for EPS isolation. This CTAB EPS isolation procedure may have caused changes in the structural properties of the EPS, potentially affecting downstream analyses. For example, a recent study by our group found that lyophilization alters the structure of cryptococcal EPS (28), and the use of detergents like CTAB have also been reported to change the physical properties of EPS (22). To examine the impact of isolation protocol on chemotype categorization we utilized a subset of cryptococcal strains expressing a single GXM motif (referred to as single motif expressing strains, SMES). These four strains are reported to express only one GXM motif – 24067 expresses M1, Mu-1 expresses M2, 409 expresses M3, and kt24066 expresses M4. Since no strains are reported to exclusively express the three remaining GXM motifs, M5, M6, and hexasaccharide, these motifs were not assessed in this analysis. For each of the four SMES, EPS was isolated using CTAB, or filtration (Supp. Figure 1). During the CTAB isolation protocol the sample was split prior to de-O-acetylation such that there were two samples, CTAB-OAc, and CTAB. This allowed us to observe the effects of the de-O-acetylation since O-acetylation is important for antibody binding {Cherniak *et al.*, 1980<u>} {</u>Belay and Cherniak, 1995}. The simplest 1D [^1^H] NMR spectra came from the CTAB sample (Figure 1, Supp. Figure 2). When comparing the CTAB and filtered EPS from the Mu-1 strain (M2 motif of GXM) the number of peaks and signal intensity was greater in the CTAB-OAc sample, and greatest in the filtered EPS sample (Figure 1). This is most notable in the SRG region (5.4 - 5.0 ppm) where the CTAB EPS contains 3 peaks while the filtered EPS contains 17 peaks. Some of the increase in complexity is due to O-acetylation as the CTAB-OAc sample contains 6 peaks in the SRG region (Supp. Figure 2). To quantify the loss of signal intensity a single culture was split, and the same volume used for CTAB and filtration preparation. Additionally, the same amount of D_6_DSS standard was added to NMR samples, allowing for quantification of signal intensity via integration of peak area. When the standard peak is set equally in each sample the loss of signal which includes O-acetylation (2.0 - 2.4 ppm) as well as within, and adjacent to, the SRG region, is clear (Figure 1). Comparison of the integrating peak area of the D_6_DSS standard to that of the SRG region shows and average ratio of 1:3.41 signal intensity between CTAB and filtered EPS (Supp. Table 4).

**Figure 1:**
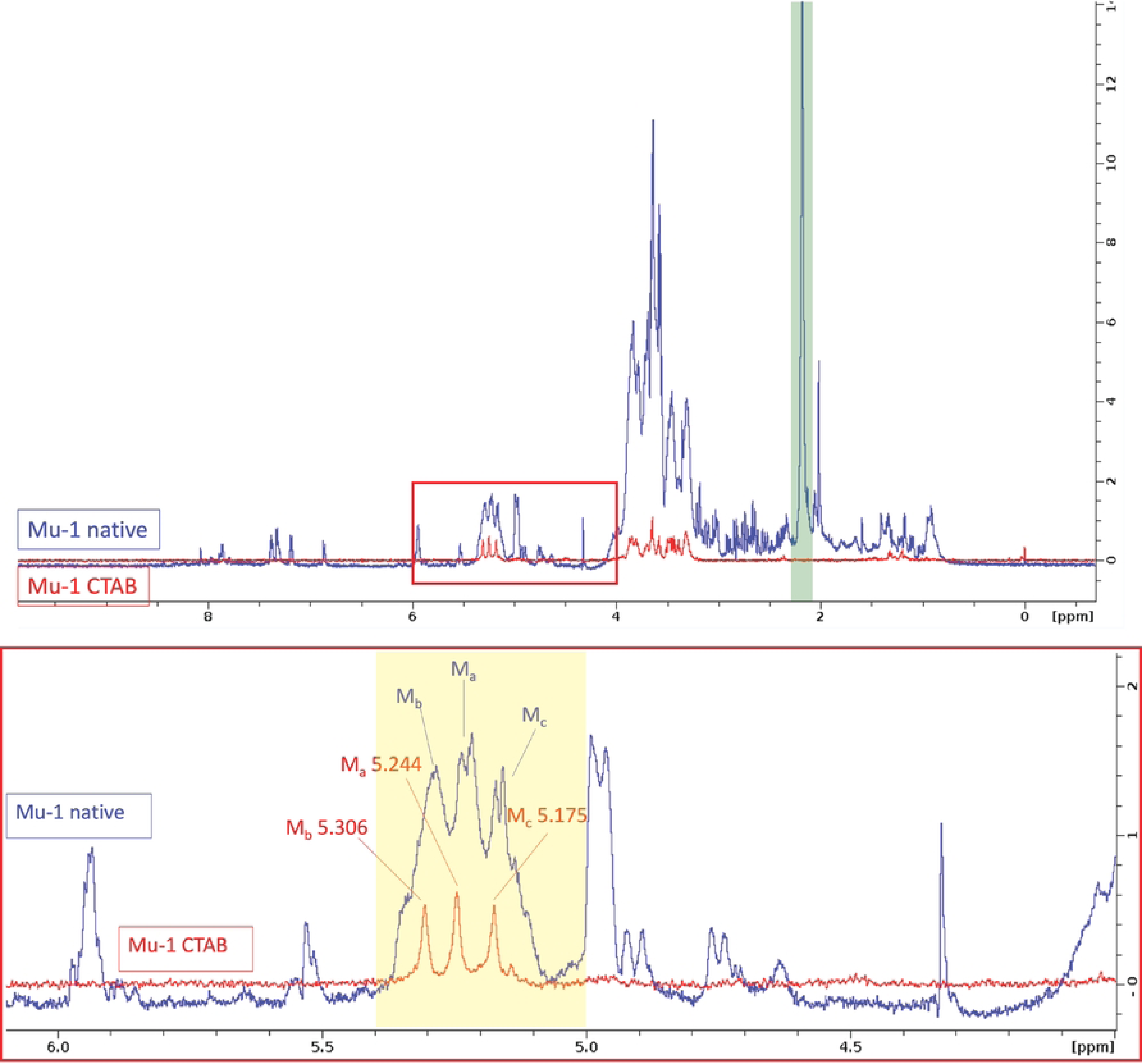
1D [^1^H] NMR analysis of Mu-1 CTAB vs. filtered EPS isolation, (top) Concentration and signal matched spectra showing signal and O-acetylation (green box) difference between samples, red boxed region peaks shown in depth (bottom) with highlighted previously reported SRG region.

### Prediction of NMR spectrum and peak chemical shift

Given differences in the observed SRG peak set between the EPS isolated by CTAB precipitation and filtration, and the limitations of 1D [^1^H] NMR, we utilized the Computer Assisted Spectrum Evaluation of Regular Polysaccharides (CASPER) computer program (http://www.casper.organ.su.se/casper/index.php) (24) to predict the proton and carbon NMR chemical shifts of the GXM M2 motif, as well as performing 2D [^1^H] [^13^C] NMR experiments on both CTAB precipitated and filtered EPS from the Mu-1 strain expressing the M2 motif. We utilized the CASPER ‘predict NMR chemical shifts’ function to simulate the 2D [^1^H] [^13^C] HSQC spectrum (24) (Figure 2a) and compared it to our own HSQC with CTAB (Figure 2b, left) and filtration isolation (Figure 2b, right) of Mu-1 EPS. With the 1D proton spectra overlaid, signals for the primary M2 motif mannose residue anomeric chemical shifts are evident in both the 1D and 2D spectra, but the 2D spectra include additional peaks between 5.0 and 4.0 ppm (Figure 2b). The CTAB preparation of Mu-1 showed no peaks 5.0 – 4.5 ppm in the 1D proton spectra (Figure 2b, left), while the filtered EPS Mu-1 sample shows a number of peaks between 5.0 – 4.5 ppm (Figure 2b, right), consistent with the notion that CTAB precipitation yielded a more homogenous preparation. In comparing the experimental NMR to the CASPER predicted spectra we observed additional differences. The CASPER data visualization does not allow for a 2D spectral view similar to those we produced from our NMR experiments, however the predicted anomeric peak chemical shifts were in the region of [^13^C] 90-110 ppm and [^1^H] 5.4-4.3 ppm. The peak at [^13^C] 100.62 ppm, [^1^H] 5.31 ppm is notably missing from the CASPER 2D HSQC visualization. This was repeatedly observed in the CASPER HSQC visualizations for GXM strain peaks above 5.0 ppm [^1^H]. Nonetheless, the CASPER predicted chemical shift table shows that the observed NMR peaks around [^1^H] 4.5 ppm represent the anomeric residues of the GlcA and Xyl residues. While the GlcA and Xyl peaks are obscured by the water peak in the 1D [^1^H] NMR, they are clear in the 2D [^1^H] [^13^C] HSQC, suggesting that in the future the SRG region of NMR could be expanded to include these peaks.

**Figure 2:**
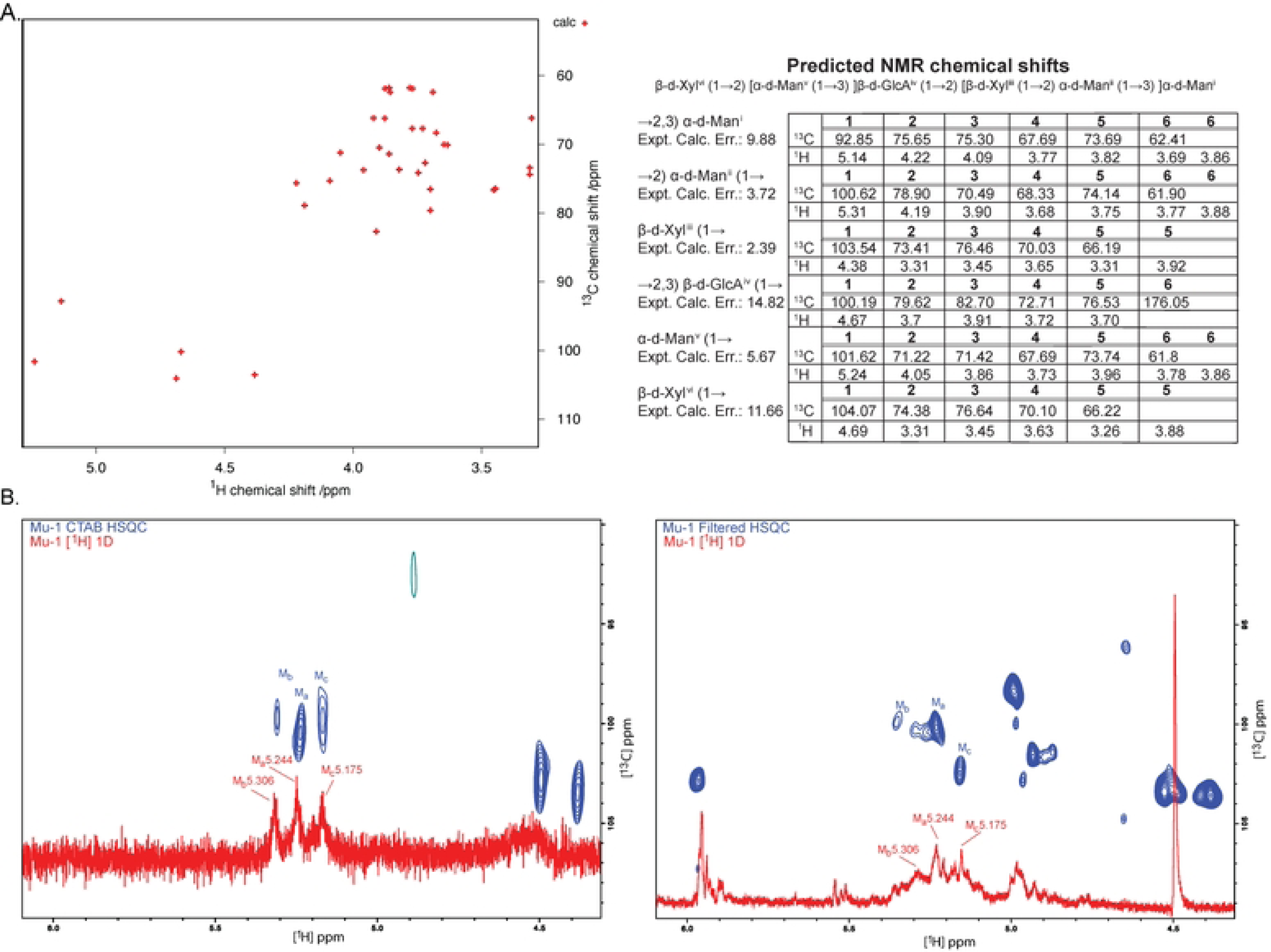
NMR and MD analysis of CTAB and physical EPS isolation. A. CASPER MD prediction of 2D [1H] (13C] HSQC peakset for the M2 motif of GXM with chemical shift table (right). B. 2D [’H] [^13^C] HSQC with 1D [H] NMR overlay of CTAB (left) and physical (right) EPS isolation.

### Consensus anomeric mannose proton chemical shifts for GXM motifs M1-M6

Since the CASPER NMR simulation and analysis of the Mu-1 samples yielded additional data about the M2 motif NMR, particularly the GlcA and Xyl residues, we were interested to extend this analysis for GXM motifs M1-M6 and compare not only the CTAB and physical isolation to the predicted chemical shifts from CASPER, but to other published NMR studies (29, 30, 31, 17) (Figure 3). We note that GXM motifs M4, M5, and M6 have far less published data than M1, M2 and M3. There are two possible reasons for this: structural complexity, as M4 and M5 are the largest and most branched GXM motifs, and lack of a strain that expresses exclusively the M5 or M6 GXM motifs. Therefore, we focused on the GXM motifs M1-M4. The CASPER predicted chemical shifts for the anomeric residues of the GXM motif saccharides showed variability from the published CTAB isolated EPS samples as well as our own filter isolated EPS samples (Figure 3b, Supp. Table 3). With the hopes of better understanding this variability and if the O-acetylation of mannose residues in GXM might play a role, we compared the recently published chemical shifts for a synthetic decasaccharide of the GXM M2 motif (32) to the CASPER predicted chemical shifts for the same structure (Supp. Table 4). Here we were able to compare the observed chemical shifts for the decasaccharide (GXM10_M2) and the de-O-acetylated version to the CASPER predicted chemical shifts for both O-acetylated and de-O-acetylated polymers. The O-acetylation of mannose did not significantly impact the chemical shifts in either the CASPER predicted, or NMR observed datasets for GXM10_M2. However, the chemical shifts of the saccharide rings varied between the CASPER predicted and NMR observed datasets (Supp. Table 4). The majority of the [^1^H] chemical shift variances were below 0.10 (44 of 68) and the majority of the [^13^C] chemical shift variance were between 0.5 and >1.0 (53 of 62) (Supp. Table 4). While the CASPER predicted chemical shifts are assigned an error, the variance observed is higher than expected, particularly in the carbon dimension. Comparison of the [^1^H] chemical shifts in (1) previously published studies, (2) CASPER prediction, and (3) our filtered EPS NMR observations showed that previously published studies and the observations of this study are more similar to each other than to the CASPER predicted chemical shifts (Supp. Table 3). This variation suggests that CASPER predicted chemical shifts may be less accurate for GXM than other published polysaccharides. However, previous published NMR chemical shifts result from cryptococcal EPS isolated using the CTAB method, which may also be hampered by the effects of the isolation method on EPS structure. We note that while our NMR observed chemical shifts for each residue showed variation of up to 0.05 ppm from previous published chemical shifts, for most residues this variation was 0.01-0.02 ppm (Figure 3b). Although we observed variations in the observed chemical shifts, this analysis did allow us to come to consensus chemical shifts for the anomeric mannose residues of each GXM motif (Figure 3b).

**Figure 3:**
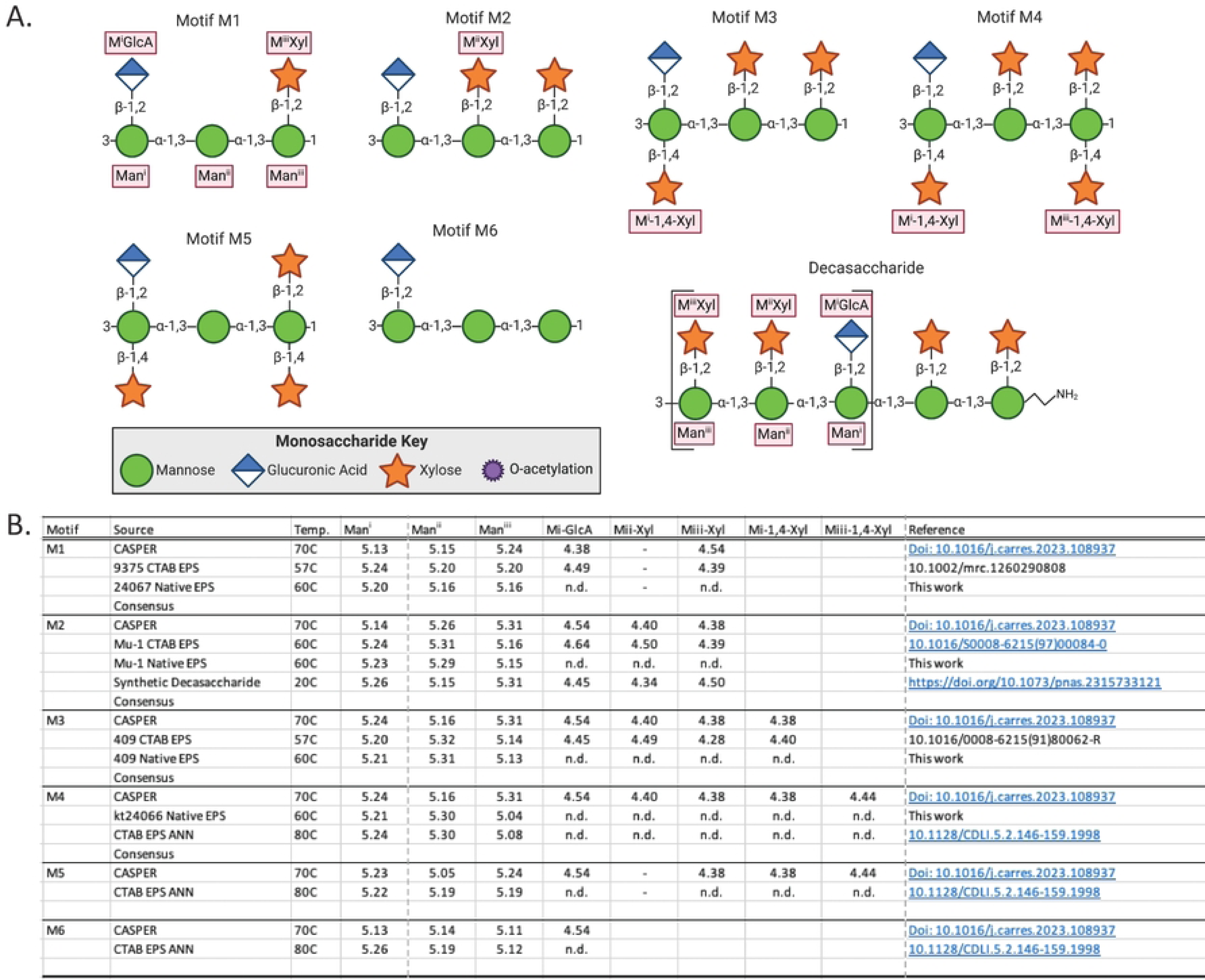
Consensus PH] chemical shifts for anomeric residues of GXM motifs. (A.) Glycobiology symbol structures of GXM motifs with residue names as used in b. (B.) Summary table of data from CASPER structure simulations, previous NMR work, artificial neural network (ANN) predicted, and current NMR data to yield consensus chemical shifts for these anomerics.

### Quantification of GXM O-acetylation in EPS

We performed peak integration analysis on 1D [^1^H] NMR spectra from a subset of strains (17) with a constant concentration of D_6_DSS added to each sample (Table 2). To assess the mannose-6-O-acetylation of these strains we set the D_6_DSS peak integral to 1 and determined the integral of the O-acetylation peak (2.20±0.05 ppm) and the SRG region (5.4 – 5.0 ppm). The ratio of the integrals of the SRG region to the O-acetylation peak we quantified as the amount of O-acetylation (Supp. Table 5). Comparison of the O-acetylation ratio with serotype, MLST genotype and chemotype showed no correlation. However, we observed a correlation between O-acetylation ratio and single motif GXM expression (Table 2). All 10 of the strains with a ratio >0.55 express a single motif GXM while 5 of the 7 strains with a ratio <0.05 express mixed motif GXM. With a sample set of only 17 strains, further study will be necessary to determine if this observation that SMES having greater O-acetylation holds.

**Table 2:**
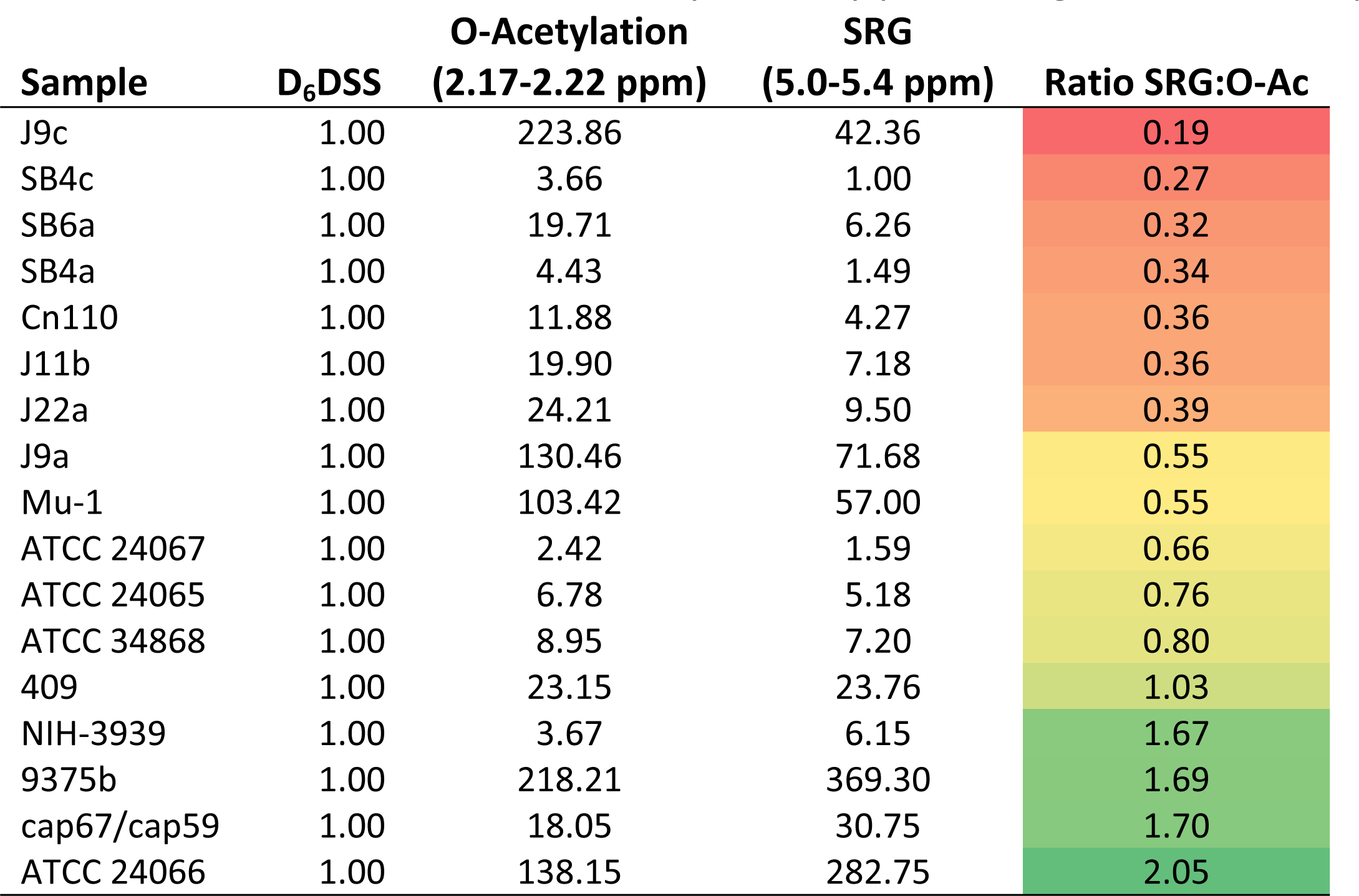
Quantification of EPS O-Acetylation by peak integration and comparison to SRG region.

### Updating Cryptococcal exopolysaccharide classification

While there are significant differences in NMR spectra between the CTAB EPS and filtered EPS, the mannose anomeric chemical shift peaks of the SRG region used in the original work defining the GXM motifs are found in both the CTAB (Figure 2b, left) and filtered EPS (Figure 2b, right) of our set of single motif expressing strains when translated from 80°C to 60°C (Supp. Figure 2). This was corroborated utilizing CASPER NMR prediction and previously published NMR studies, from which we have established a consensus set of proton chemical shifts for GXM motif identification at 60°C (Figure 3). Throughout this analysis we noted that the chemotyping system of GXM motif categorization was often not descriptive of the motif expressed. Along with the fact that filtered EPS 1D [^1^H] NMR showed chemical shift peaks outside of the original CTAB EPS reported SRG region, our analysis showed that there is no correlation between chemotype and genotype, with the caveat that MLST does not include genes shown to be directly involved in capsular construction. Therefore, we felt it was important to update the system by which we classified cryptococcal strain GXM motif expression in EPS. First, we recommend the simpler EPS isolation by sterile filtration technique over CTAB isolation. Second, a facile classification system can be developed based on GXM motif expression. For this classification there are three types, (i) SMES (ii) mixed motif strains, and (iii) mixed motif dominant strains (Figure 4). If a strain expresses a single motif, then it can be identified by that motif. For example, Mu-1 expresses exclusively the M2 motif and is thus a M2 motif-expressing strain and could be referred to as a “M2 SMES”. An alternative situation, in which strains express more than one motif where no lone motif is dominant, we propose the classification “mixed motif strain”. For instance, the strain 9375b expresses M1, M5 and M6, so it is a mixed motif strain. A special case of the mixed motif strain arises when one motif represents the majority of the GXM composition. Again, let us use the strain 9375b which expresses three GXM motifs but is 74% M1. This strain would be referred to as a “mixed motif, M1 strain” to denote this motif makes up more than 50% of the motif expression. In the future, we hope to introduce O-acetylation into cryptococcal classification, but this will require more data to confirm that SMES have greater O-acetylation than mixed motif strains. To facilitate the adoption of PS structure into cryptococcal strain classification we have put together a simple workflow (Supp. Figure 1). This workflow introduces the physical EPS isolation technique utilized throughout this work which is less intensive than CTAB isolation. Further, this method allows for PS classification of strains with significantly reduced material (10 mL culture) and time commitment (2 days).

**Figure 4:**
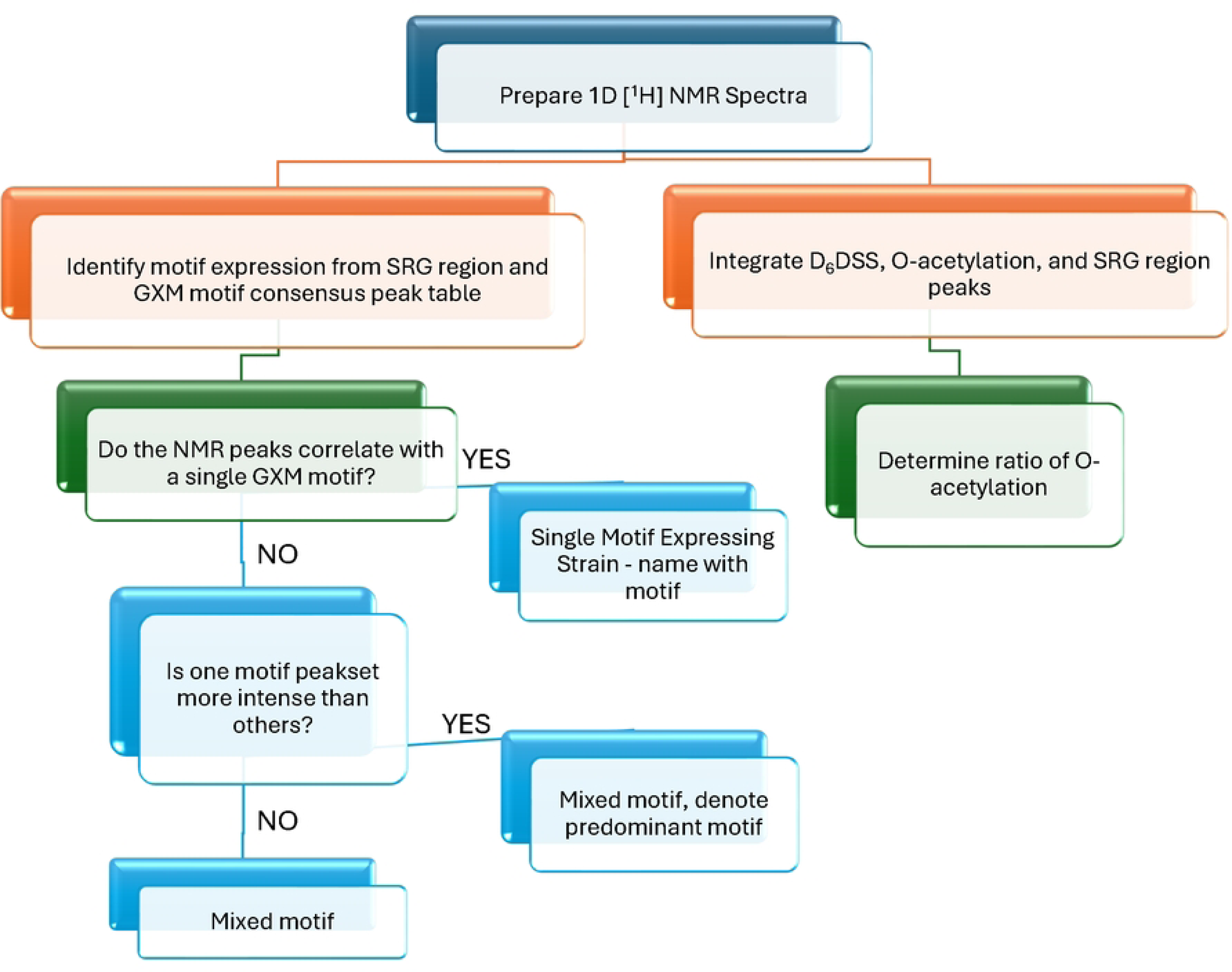

## Discussion

Through this work we were surprised to find that while there is a weak correlation between serotype and MLST classifications, we were unable to correlate serotype and chemotype or GXM motif expression. Hence, the often-assumed conclusion that differences in polysaccharide structure translate into antigenic differences that confer serotype classification is not tenable. While serotype A strains are reported to express the M2 motif and should be chemotype 4, these strains include chemotypes 1, 2, 3, 4, and 5 and express all GXM motifs save M4. A deep dive into the literature reveals that serotype is not, in fact, related to GXM motif (9, 10, 11, 12). Serotype reflects a classification obtained by antibody reactivity with an antigen on the surface of the cryptococcal capsule that, when bound by bivalent antibodies, such as those found in sera, results in yeast cell agglutination. A limitation of serotype classification is that polyclonal antibody reagents obtained from small animals are not constant as they depend on method of immunization, the type of host and the immunological state of the host. Currently, those reagents are no longer available, and the nature of this agglutinating antigen is unknown. Knowing that the capsule contains more than just polysaccharides, it is likely that the adsorbed anti-sera initially utilized to determine serotype targeted a factor within the capsular antigen that is different from GXM, possibly a different polysaccharide or protein. Given that capsules contain lipids (33)(34) it is even conceivable that the serotype-conferring antigen is a lipid, lipoprotein or lipopolysaccharide. Since neither serotype nor MLST classification include the contribution of polysaccharide structure, we were curious if GXM motif expression aligned with MLST.

In our analysis of MLST and GXM motif expression we first consider the chemotyping system. In our table of 63 strains, 18 strains are VNI, the chemotypes include 4 strains of chemotype 2, 6 strains of chemotype 3, and 5 strains of chemotype 4, and 4 strains of chemotype 5. The GXM motif expression is more expansive than the chemotypes, including only two strains that are 100% M2 expressing, but 4 strains expressing >45% M1, 6 strains expressing >45% M2, and 7 strains expressing 3 or more GXM motifs with none at >45% of the total. This is a high level of variability within a single genotype with no apparent conserved GXM motif. Additionally, our analysis revealed several strains that, while attributed with distinct numbers, tracked back to the same strain. Together this suggests that MLST genotyping, while useful for strain categorization, does not account for GXM motif expression and therefore excludes this essential virulence factor. At this juncture the genetic regulation of GXM synthesis is unclear, along with any regulation of GXM motif expression.

To better understand GXM motif expression across strains and determine a method of classification to accompany MLST genotyping, we returned to literature information. Seminal work done by Cherniak and colleagues and Battacharjee and colleagues lead to the characterization of repeat units for both GXM and GXMgal (17, 35, 36, 15, 29, 37, 38, 39, 40, 41, 42). While GXMgal is an important component, it is also minor, making up just 3-10% of EPS (17, 35, 36, 15, 29, 37, 38, 39, 40, 41, 42). Constituting up to 90% of cryptococcal polysaccharides, GXM has long been the major glycan of interest. When techniques for the isolation of PS were initially developed CTAB precipitation was found to be an effective method for separating GXM from GXMgal. However, CTAB is a detergent and difficult to remove from the PS sample, requiring days of dialysis to purify samples, and one could never be certain that all was removed. Additionally, the lack of sterile conditions in dialyzing out the CTAB required lyophilization of samples for long-term storage, which we now know introduced additional structural changes (28). Previous work by our group found that isolation of EPS from culture supernatant by filtration was possible (22). Though that work fractionated the EPS by molecular mass and used filtration that was not sterile, we were able to update the methodology using sterile filtration (28). However, we were uncertain if these samples would yield the same results with regard to GXM motif as the original CTAB preparations. Fortunately, the SRG peak sets originally reported for GXM motifs were retained in both samples, providing consistency and reproducibility among these methods. To examine the SRG chemical shifts across as many datasets as possible for the SMES we coupled our NMR data with that of published papers as well as CASPER simulation predicted 2D [^1^H] [^13^C] chemical shifts of the GXM motifs (24).

Together these chemical shifts showed variation, likely due to biological variables we have yet to identify, but within an acceptable tolerance to establish consensus chemical shifts within the SRG region for each GXM motif. Since the previously established chemotypes do not align with any other strain characterization, we simplified the categorization method to three types (i) SMES (ii) strains expressing multiple motifs but predominantly (>50%) one GXM motif and (iii) strains expressing two or more GXM motifs with no dominant motif. Within the 63-strain collection examined, 18 were SMES while 45 were mixed motif, of which 23 expressed a dominant motif. Unsurprisingly, this updated GXM motif classification still does not show alignment with MLST genotyping. However, our analysis of a subset of strains shows that high levels of mannose-6-O-acetylation, which is necessary for antibody binding, correlate with the expression of a single GXM motif. Additionally, this method does not focus on the four GXM motifs originally assigned to serotypes but allows for the contribution of all six GXM motifs to strain classification.

The lack of association between genotype and chemotype, or GXM motif expression in EPS challenges the way we have perceived the association between genotype and capsule type. Work in encapsulated bacteria suggests that a set of genes are responsible for capsular and exo-polysaccharide expression and that these genes can be linked to serotype by patterns of antigen immunogenicity and agglutination (43, 44). However, in this analysis we observe three sets of strains with the same origin, yet dramatically different GXM motif expression. While this has previously been explored for the strain 24067 and termed GXM microheterogeneity (27), our analysis shows greater variance than described for 24067.

The listing for strain ATCC 28958 shows another strain designation of 430, which was described as a distinct strain in the original chemotyping work (17). Additionally, while both strains expressed GXM motifs M1, M5 and M6, their reported percentages differed. While 430 is 38% M1, 44% M5, and 18% M6, ATCC 28958 is 34% M1, 19% M5, and 47% M6. Together with the GXM variance described for 24067, this suggests that in *Cryptococcus* the regulation of GXM motif expression may have a non-genetically inherited component. This observation requires future research investigating epigenetic and other non-genetic mechanisms of inheritance.

In addition to the complexity described here for the relationships between serotype, chemotype and genotype, we note that both EPS structure and genetic changes can occur during animal and human passage. Analysis of serial isolates recovered from individual patients belonging to the same genotype revealed differences in EPS structure (45). Phenotypic switching of a single strain can result in changes in the proportion of GXM reporter groups (46). Comparison of genomic sequence of a strain recovered from a pet cockatoo that was implicated in a point source infection with those recovered from a patient and after mouse passage revealed genomic changes occurring during mammalian infection (47). Hence, any classification scheme for *Cryptococcus* spp. must contend with the fact that isolates are highly malleable with regards to genotype and phenotype.

Beyond the questions and uncertainties that this work has unraveled, it has also yielded new insights into difficulties involving vaccine development. For instance, the lack of association between serotype and chemotype suggests that oligosaccharide vaccines synthesized based on GXM motifs can be expected to elicit very different antibody responses, than those generated in response to whole cells. Further, this could explain the disappointing efficacy of synthetic oligosaccharide conjugate vaccines (48, 49, 50). Additionally, the search for genes regulating cryptococcal polysaccharide synthesis has proved far more difficult than other cryptococcal pathways (51). The lack of association between genotype and chemotype or GXM motif expression suggests that the polysaccharide synthesis pathway may be more complex than previously hypothesized and may involve non-genetic regulation. The role of metabolism in producing chemotype diversity remains unexplored. As technologies continue to advance, one can expect to add more details to the analysis of EPS, GXM motifs, and cryptococcal polysaccharide structure. Hopefully it will be possible to add genes and perhaps epigenetic regulators to MLST strain analysis such that molecular type will also reflect GXM motif expression. Until this is possible, we hope that the rapid and facile method proposed here will enable researchers to include classification of this critical virulence factor into cryptococcal strain descriptions.

---

**Supp Fig 1:**
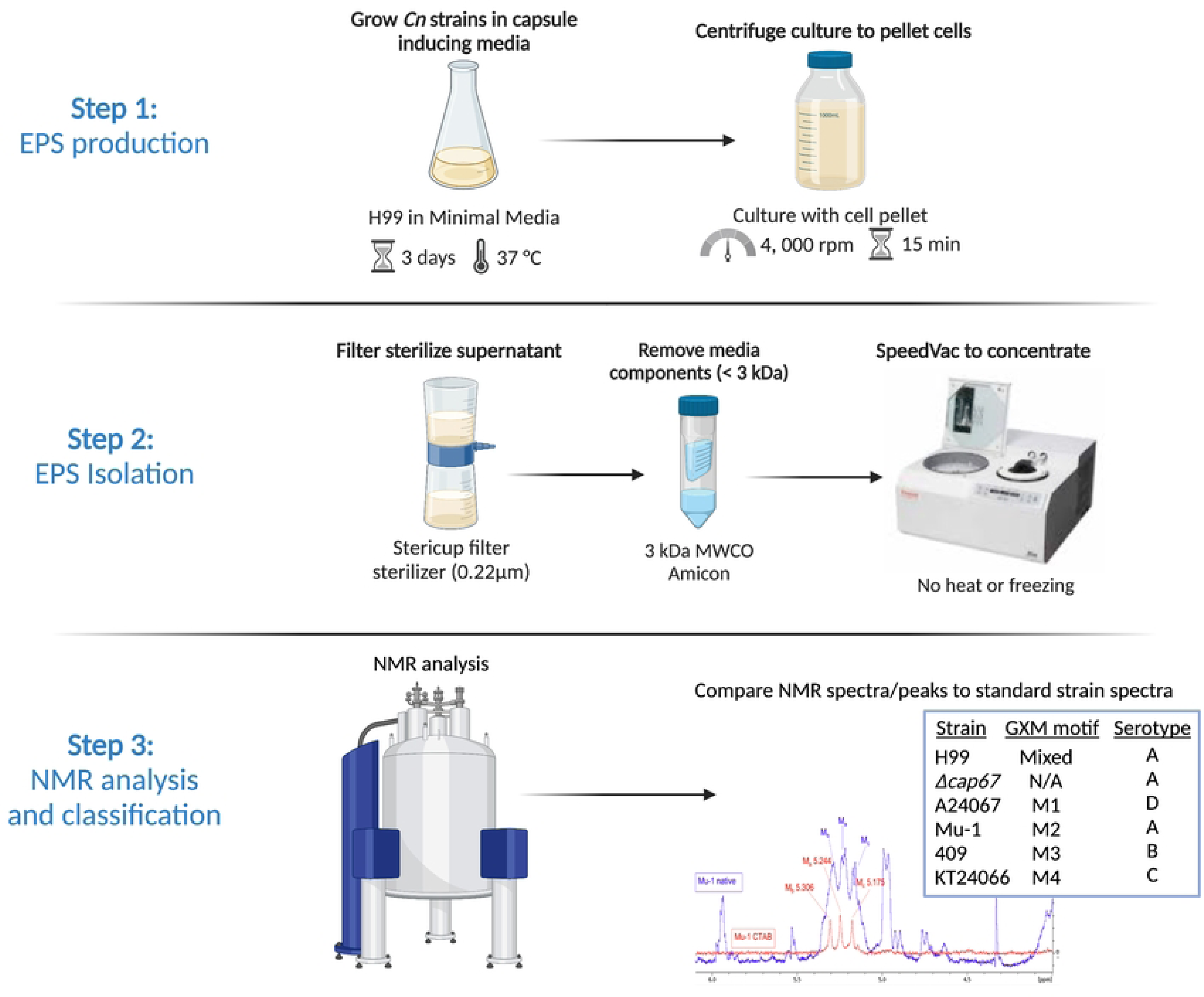
Workflow for updated EPS classification. Three step process to classify EPS from a cryptococcal strain. This includes culture, PS isolation, and NMR analysis for updated classification.

**Supp Fig 3:** 

**Supp Fig 4:**
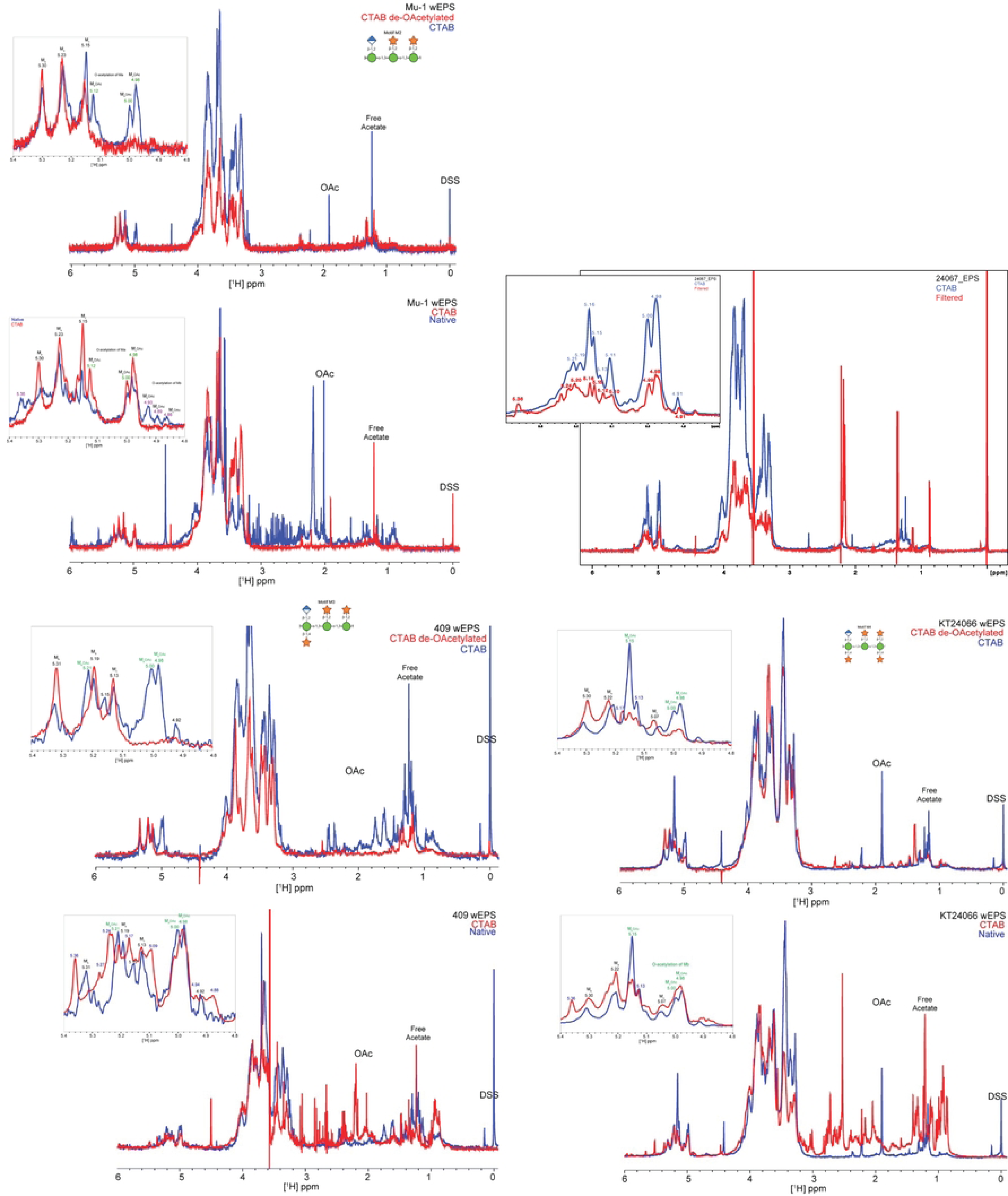

## Bibliography

1. Kwon-Chung KJ, Bennett JE, Wickes BL, Meyer W, Cuomo CA, Wollenburg KR, et al. The case for adopting the “species complex” nomenclature for the etiologic agents of cryptococcosis. mSphere. 2017 Jan 11;2(1).

2. McClelland EE, Bernhardt P, Casadevall A. Estimating the relative contributions of virulence factors for pathogenic microbes. Infect Immun. 2006 Mar;74(3):1500–4.

3. Zaragoza O, Rodrigues ML, De Jesus M, Frases S, Dadachova E, Casadevall A. Chapter 4 The Capsule of the Fungal Pathogen Cryptococcus neoformans. Elsevier; 2009. p. 133–216.

4. Albuquerque PC, Fonseca FL, Dutra FF, Bozza MT, Frases S, Casadevall A, et al. Cryptococcus neoformans glucuronoxylomannan fractions of different molecular masses are functionally distinct. Future Microbiol. 2014;9(2):147–61.

5. Araujo G de S, Fonseca FL, Pontes B, Torres A, Cordero RJB, Zancopé-Oliveira RM, et al. Capsules from pathogenic and non-pathogenic Cryptococcus spp. manifest significant differences in structure and ability to protect against phagocytic cells. PLoS One. 2012 Jan 12;7(1):e29561.

6. Grechi J, Marinho-Carvalho M, Zancan P, Cinelli LP, Gomes AMO, Rodrigues ML, et al. Glucuronoxylomannan from Cryptococcus neoformans down-regulates the enzyme 6-phosphofructo-1-kinase of macrophages. J Biol Chem. 2011 Apr 29;286(17):14820–9.

7. Barbosa FM, Fonseca FL, Figueiredo RT, Bozza MT, Casadevall A, Nimrichter L, et al. Binding of glucuronoxylomannan to the CD14 receptor in human A549 alveolar cells induces interleukin-8 production. Clin Vaccine Immunol. 2007 Jan;14(1):94–8.

8. Vecchiarelli A, Pericolini E, Gabrielli E, Kenno S, Perito S, Cenci E, et al. Elucidating the immunological function of the Cryptococcus neoformans capsule. Future Microbiol. 2013 Sep;8(9):1107–16.

9. BENHAM RW. Cryptococci: Their Identification by Morphology and by Serology on JSTOR. The Journal of Infectious Diseases,. 1935 Nov;Vol. 57(No. 3):255–74.

10. Evans EE, Kessel JF. The antigenic composition of cryptococcus neoformans. The Journal of Immunology. 1951 Aug 1;67(2):109–14.

11. Evans EE. The antigenic composition of cryptococcus neoformans. The Journal of Immunology. 1950 May 1;64(5):423–30.

12. Wilson DE, Bennett JE, Bailey JW. Serologic grouping of Cryptococcus neoformans. Proc Soc Exp Biol Med. 1968 Mar;127(3):820–3.

13. Bennett JE, Kwon-Chung KJ, Howard DH. Epidemiologic differences among serotypes of Cryptococcus neoformans. Am J Epidemiol. 1977 Jun;105(6):582–6.

14. Mishra SK, Staib F, Folkens U, Fromtling RA. Serotypes of Cryptococcus neoformans strains isolated in Germany. J Clin Microbiol. 1981 Jul;14(1):106–7.

15. Heiss C, Klutts JS, Wang Z, Doering TL, Azadi P. The structure of Cryptococcus neoformans galactoxylomannan contains beta-D-glucuronic acid. Carbohydr Res. 2009 May 12;344(7):915–20.

16. Bhattacharjee AK, Kwon-Chung KJ, Glaudemans CPJ. On the structure of the capsular polysaccharide fromCryptococcus neoformans serotype C. Immunochemistry. 1978 Aug;15(9):673–9.

17. Cherniak R, Valafar H, Morris LC, Valafar F. Cryptococcus neoformans chemotyping by quantitative analysis of 1H nuclear magnetic resonance spectra of glucuronoxylomannans with a computer-simulated artificial neural network. Clin Diagn Lab Immunol. 1998 Mar;5(2):146–59.

18. Brandt S, Thorkildson P, Kozel TR. Monoclonal antibodies reactive with immunorecessive epitopes of glucuronoxylomannan, the major capsular polysaccharide of Cryptococcus neoformans. Clin Diagn Lab Immunol. 2003 Sep;10(5):903–9.

19. Litvintseva AP, Thakur R, Vilgalys R, Mitchell TG. Multilocus sequence typing reveals three genetic subpopulations of Cryptococcus neoformans var. grubii (serotype A), including a unique population in Botswana. Genetics. 2006 Apr;172(4):2223–38.

20. Meyer W, Aanensen DM, Boekhout T, Cogliati M, Diaz MR, Esposto MC, et al. Consensus multi-locus sequence typing scheme for Cryptococcus neoformans and Cryptococcus gattii. Med Mycol. 2009;47(6):561–70.

21. Nimrichter L, Frases S, Cinelli LP, Viana NB, Nakouzi A, Travassos LR, et al. Self-aggregation of Cryptococcus neoformans capsular glucuronoxylomannan is dependent on divalent cations. Eukaryotic Cell. 2007 Aug;6(8):1400–10.

22. Frases S, Nimrichter L, Viana NB, Nakouzi A, Casadevall A. Cryptococcus neoformans capsular polysaccharide and exopolysaccharide fractions manifest physical, chemical, and antigenic differences. Eukaryotic Cell. 2008 Feb;7(2):319–27.

23. Cordero RJB, Frases S, Guimaräes AJ, Rivera J, Casadevall A. Evidence for branching in cryptococcal capsular polysaccharides and consequences on its biological activity. Mol Microbiol. 2011 Feb;79(4):1101–17.

24. Dorst KM, Widmalm G. NMR chemical shift prediction and structural elucidation of linker-containing oligo- and polysaccharides using the computer program CASPER. Carbohydr Res. 2023 Nov;533:108937.

25. Kenosi K, Mosimanegape J, Daniel L, Ishmael K. Recent advances in the ecoepidemiology, virulence and diagnosis of cryptococcus neoformans and cryptococcus gattii species complexes. Open Microbiol J. 2023 Apr 19;17(1).

26. Hagen F, Khayhan K, Theelen B, Kolecka A, Polacheck I, Sionov E, et al. Recognition of seven species in the Cryptococcus gattii/Cryptococcus neoformans species complex. Fungal Genet Biol. 2015 May;78:16–48.

27. Franzot SP, Mukherjee J, Cherniak R, Chen LC, Hamdan JS, Casadevall A. Microevolution of a standard strain of Cryptococcus neoformans resulting in differences in virulence and other phenotypes. Infect Immun. 1998 Jan;66(1):89–97.

28. Wear MP, Hargett AA, Kelly JE, McConnell SA, Crawford CJ, Freedberg DI, et al. Lyophilization induces physicochemical alterations in cryptococcal exopolysaccharide. Carbohydr Polym. 2022 Sep 1;291:119547.

29. Skelton MA, Cherniak R, Poppe L, van Halbeek H. Structure of the De-O-acetylated glucuronoxylomannan fromCryptococcus neoformans serotype D, as determined by 2D NMR spectroscopy. Magn Reson Chem. 1991 Aug;29(8):786–93.

30. Sheng S, Cherniak R. Structure of the 13C-enriched O-deacetylated glucuronoxylomannan of Cryptococcus neoformans serotype A determined by NMR spectroscopy. Carbohydr Res. 1997 Jun;301(1–2):33–40.

31. Hargett AA, Azurmendi HF, Crawford CJ, Wear MP, Oscarson S, Casadevall A, et al. The structure of a C. neoformans polysaccharide motif recognized by protective antibodies: A combined NMR and MD study. Proc Natl Acad Sci USA. 2024 Feb 13;121(7):e2315733121.

32. Hargett AA, Azurmendi HF, Crawford CJ, Wear MP, Oscarson S, Casadevall A, et al. The structure of a C. neoformans polysaccharide motif recognized by protective antibodies: A combined NMR and MD study. BioRxiv. 2023 Sep 6;

33. Nicola AM, Frases S, Casadevall A. Lipophilic dye staining of Cryptococcus neoformans extracellular vesicles and capsule. Eukaryotic Cell. 2009 Sep;8(9):1373–80.

34. Sebolai OM, Pohl CH, Botes PJ, Strauss CJ, van Wyk PWJ, Botha A, et al. 3-hydroxy fatty acids found in capsules of Cryptococcus neoformans. Can J Microbiol. 2007 Jun;53(6):809–12.

35. Bhattacharjee AK, Kwon-Chung KJ, Glaudemans CP. Capsular polysaccharides from a parent strain and from a possible, mutant strain of Cryptococcus neoformans serotype A. Carbohydr Res. 1981 Sep 16;95(2):237–48.

36. Cherniak R, O’Neill EB, Sheng S. Assimilation of xylose, mannose, and mannitol for synthesis of glucuronoxylomannan of Cryptococcus neoformans determined by 13C nuclear magnetic resonance spectroscopy. Infect Immun. 1998 Jun;66(6):2996–8.

37. Turner SH, Cherniak R, Reiss E, Kwon-Chung KJ. Structural variability in the glucuronoxylomannan of Cryptococcus neoformans serotype A isolates determined by 13C NMR spectroscopy. Carbohydr Res. 1992 Sep 2;233:205–18.

38. Cherniak R, Morris LC, Turner SH. Glucuronoxylomannan of Cryptococcus neoformans serotype D: structural analysis by gas-liquid chromatography-mass spectrometry and by 13C-nuclear magnetic resonance spectroscopy. Carbohydr Res. 1992 Jan;223:263–9.

39. Cherniak R, Morris LC, Meyer SA. Glucuronoxylomannan of Cryptococcus neoformans serotype C: structural analysis by gas-liquid chromatography-mass spectrometry and 13C-nuclear magnetic resonance spectroscopy. Carbohydr Res. 1992 Mar 2;225(2):331–7.

40. Turner SH, Cherniak R, Reiss E. Fractionation and characterization of galactoxylomannan from Cryptococcus neoformans. Carbohydr Res. 1984 Feb 15;125(2):343–9.

41. Skelton MA, van Halbeek H, Cherniak R. Complete assignment of the 1H- and 13C-n.m.r. spectra of the O-deacetylated glucuronoxylomannan from Cryptococcus neoformans serotype B. Carbohydr Res. 1991 Dec 16;221:259–68.

42. Turner SH, Cherniak R. Glucuronoxylomannan of Cryptococcus neoformans serotype B: structural analysis by gas-liquid chromatography-mass spectrometry and 13C-nuclear magnetic resonance spectroscopy. Carbohydr Res. 1991 Apr 2;211(1):103–16.

43. Weintraub A. Immunology of bacterial polysaccharide antigens. Carbohydr Res. 2003 Nov 14;338(23):2539–47.

44. Weinberger DM, Trzciński K, Lu Y-J, Bogaert D, Brandes A, Galagan J, et al. Pneumococcal capsular polysaccharide structure predicts serotype prevalence. PLoS Pathog. 2009 Jun 12;5(6):e1000476.

45. Cherniak R, Morris LC, Belay T, Spitzer ED, Casadevall A. Variation in the structure of glucuronoxylomannan in isolates from patients with recurrent cryptococcal meningitis. Infect Immun. 1995 May;63(5):1899–905.

46. Fries BC, Goldman DL, Cherniak R, Ju R, Casadevall A. Phenotypic switching in Cryptococcus neoformans results in changes in cellular morphology and glucuronoxylomannan structure. Infect Immun. 1999 Nov;67(11):6076–83.

47. Sephton-Clark P, McConnell SA, Grossman N, Baker RP, Dragotakes Q, Fan Y, et al. Similar evolutionary trajectories in an environmental Cryptococcus neoformans isolate after human and murine infection. Proc Natl Acad Sci USA. 2023 Jan 10;120(2):e2217111120.

48. Crawford CJ, Liporagi-Lopes L, Coelho C, Santos Junior SR, Nicola AM, Wear MP, et al. Semi-synthetic glycoconjugate vaccine candidate against Cryptococcus neoformans. BioRxiv. 2024 Feb 3;

49. Nakouzi A, Zhang T, Oscarson S, Casadevall A. The common Cryptococcus neoformans glucuronoxylomannan M2 motif elicits non-protective antibodies. Vaccine. 2009 Jun 2;27(27):3513–8.

50. Oscarson S, Alpe M, Svahnberg P, Nakouzi A, Casadevall A. Synthesis and immunological studies of glycoconjugates of Cryptococcus neoformans capsular glucuronoxylomannan oligosaccharide structures. Vaccine. 2005 Jun 10;23(30):3961–72.

51. O’Meara TR, Alspaugh JA. The Cryptococcus neoformans capsule: a sword and a shield. Clin Microbiol Rev. 2012 Jul;25(3):387–408.

